# Prevalence of UGT1A1 Genetic Variants in Argentinean Population, Potential Implications for Pharmacogenomic Testing

**DOI:** 10.1101/768994

**Authors:** Cintia V. Cruz, Paula Scibona, Waldo H. Belloso

## Abstract

The four groups of uridine diphosphate glucuronosyltransferase (UGT) enzymes form a superfamily responsible for the glucuronidation of target substrates. These include hormones, flavonoids, environmental mutagens and pharmaceutical drugs. Thus, UGT enzymes are relevant for pharmacogenetic research.

Most of the members of the UGT family are expressed in the liver, but are also present in intestinal, stomach or breast tissue.

The incidences and types of polymorphisms for different enzymes vary with geographical regions and ethnic groups. This is the first study that examined the frequency of polymorphisms for UGT1 isozymes for a population of 100 healthy argentinians.

The distribution of UGT1A1 in our population was: 70.5% (70.5) for the *1 allele, 21.5% for the *28 allele and 1% for the *36 allele. 48% (48) presented the *1/*1 genotype, while 43 % (43) had *1/*28, 2% (2) had *1/*36 and 7% (7) showed *28/*28. There was no preferential sex distribution.

Since most Argentinians are of Caucasian descent, a European genotype frequency profile is to be expected. That is evident in the wild type prevalence in our population. However, the contribution of Native American ancestry to gene pool components may in part explain the higher prevalence of the *28 genotype in UGT1A1 *1 in our population, in comparison with European cohorts.

## INTRODUCTION

The uridine diphosphate glucuronosyltransferase (UGT) enzymes conform a superfamily of enzymes responsible for the glucuronidation of target substrates. The transfer of glucuronic acid renders xenobiotics and other endogenous compounds water soluble, allowing for their biliary or renal excretion. The UGT superfamily is responsible for the glucuronidation of hundreds of compounds, including hormones, flavonoids and environmental mutagens. Most of the members of the UGT family are expressed in the liver, as well as in other type of tissues, such as intestinal, stomach or breast. A few members are expressed only outside the liver such as UGT1A7, UGT1A8, UGT1A10 and UGT2A1. UGT superfamily is composed by four families: UGT1A, UGT2, UGT3 and UGT8. UGT2 is further divided into two subfamilies, UGT2A and UGT2B, both of which present on chromosome 4. Although limited studies are already available on UGT2A enzymes, they appear to be involved in the glucuronidation of compounds such as phenolic odorants and polycyclic aromatic hydrocarbon metabolites. UGT2B proteins are mainly responsible for the metabolism of steroids. The roles of UGT3 and UGT8 family members have not been well characterized yet.

The UGT1A family is located on chromosome 2q37(1), and members of this group glucuronidate a large variety of compounds. Pharmaceutical drugs are also a common substrate of the UGT family, turning the enzymes in this group relevant for pharmacogenetic research(2). Bilirubin-UGT (*UGT1A1*) conjugates bilirubin with glucuronic acid, converting the bilirubin into a water-soluble form that is readily excreted in bile. Mutations of bilirubin UDP-glucurunosyl transferase causes hereditary unconjugated hyperbilirubinemias, including Crigler-Najjar and Gilbert syndromes(3).

The genetic defect in patients with Gilbert syndrome involves the promoter region of *UGT1A1*(4,5). Gilbert syndrome manifests only in people homozygous for the variant promoter. As a result, its inheritance is most consistent with an autosomal recessive trait. However, heterozygotes for the Gilbert genotype have higher average plasma bilirubin concentrations compared with those with two wild-type alleles. It is estimated that 9 percent of individuals in the Western general population world are homozygous for the variant promoter, and up to 42 percent are heterozygous(5).

Characterization of the *UGT1A* gene locus has permitted an understanding of the molecular defects responsible for the Gilbert syndrome. The mutation responsible for Gilbert syndrome is in the promoter region, upstream to exon 1 of *UGT1A1*. The normal sequence of the TATAA element within the promoter is A[TA]6TAA. Caucasian and black patients with Gilbert syndrome are homozygous for a longer version of the TATAA sequence, A[TA]7TAA, which causes reduced production of bilirubin-UGT. This variant is termed *UGT1A1*28*(6).

This longer TATAA element has been found in all individuals with Gilbert syndrome studied in the United States, Europe, and countries of the Middle East and South Asia. However, other factors are probably involved in the expression of Gilbert phenotype since not all patients who are homozygous for the variant promoter develop hyperbilirubinemia. Furthermore, in the Japanese population, other mutations within the coding regions of *UGT1A1* can be the underlying cause of the Gilbert phenotype.

Since bilirubin-UGT is involved in the glucuronidation of several important drugs, individuals with Gilbert syndrome may be more susceptible to the toxic effect of substances that require bilirubin-UGT-mediated hepatic glucuronidation prior to excretion. Gilbert syndrome is known to increase the risk of drug toxicity with irinotecan and the hyperbilirubinemia associated with atazanavir(2,7).

The active metabolite of irinotecan, SN-38, is glucuronized in the liver mainly by bilirubin-UGT. The major dose-limiting toxicity of this drug is diarrhoea.In patients who inherit certain *UGT1A1* polymorphisms, reduced glucuronidation of SN-38 leads to an increased incidence of diarrhoea. The symptoms can be severe enough to warrant switching to other drugs. Thus, it is currently recommended to test the UGT1A1 polymorphism prior to the administration of Irinotecan and to adjust the dose according to the resulting genotype(8).

As previously stated, some drugs may induce hyperbilirubinemia in patients with Gilbert syndrome. Atazanavir, an antiretroviral medication, is an inhibitor of bilirubin-UGT activity and is associated with hyperbilirubinemia(9). Isolated hyperbilirubinemia has also been reported during the treatment of hepatitis C with peginterferon and ribavirin and in patients receiving pazopanib. In such cases, discontinuation of therapy is usually not necessary(10).

The incidences and types of the polymorphisms for these enzymes are quite different according to geographical regions and ethnic groups. The aim of this study is to estimate the prevalence of UGT1A1 polymorphism in the Argentine population and to evaluate what clinical implications this might have.

## MATERIAL AND METHODS

<u>dx.doi.org/10.17504/protocols.io.6sjhecn</u>

### Study population

One hundred random and anonymized DNA samples from healthy donors were analysed. The Hospital Italiano de Buenos Aires DNA Bank collection project has the approval of the local ethic committee and all the volunteer subjects signed an informed consent.

### Sample collection and DNA extraction

Following an informed consent process, 10 mL of peripheral blood were collected from each subject in 5-mL EDTA tubes. Whole blood samples were stored at 4ºC until the time of processing. Genomic DNA was extracted and purified using the QI Amp DNA Blood Mini kit (QIAGEN).

### Genotyping of UGT1A1

Genomic DNA was extracted from 200 μL of whole blood using QIAamp DNA Mini kit (Qiagen, GmbH, D-40724 Hilden, Germany). The polymerase chain reaction (PCR) was performed in a final volume of 20 μL. The forward primer was 5’-CAGCCTCAAGACCCCACA – 3’ and the reverse primer was 5’-TGCTCCTGCCAGAGGTTC -3’. The PCR conditions were 5 minutes at 95°C, followed by 35 cycles of 30 seconds at 95°C, 60 seconds at 61°C, 60 seconds at 72°C, and final extension for 10 minutes at 72°C. The PCR product was detected on 2% agarose gels by means of ethidium bromide staining. The presence of variant UGT1A1 was confirmed by direct sequencing of PCR products on an automated ABI 3100 capillary sequencer (Applied Biosystems, Foster City, Calif) using the Big Dye Terminator Cycle Sequencing Kit (Applied Biosystems).

## STATISTICAL ANALYSIS

Allelic and genotypic frequencies were expressed in absolute and percentage values in the study population. Sample size calculation based on the estimation of a 4% prevalence of TT genotype as published for european population was of 64 samples, while for a prevalence of 7 % as in global population was of 110 samples, with an alpha error of 5 %. The results obtained were further evaluated for Hardy-Weinberg equilibrium.

## RESULTS

The genotyping of all samples was according to the method previously described. Among the 100 subjects analysed, 49% (49) were male and 51% (51) were female with a median age of 43 and a range of 32-80 years.

In our population, the distribution of UGT1A1 was -Figure 1 attached separatedly-: 70.5% (70.5) for the *1 allele, 21.5% for the *28 allele and 1% for the *36 allele. 48% (48) presented the *1/*1 genotype, while 43 % (43) had *1/*28, 2% (2) had *1/*36 and 7% (7) showed *28/*28. No preferential sex distribution was observed between genotypes.

**Figure 1:**
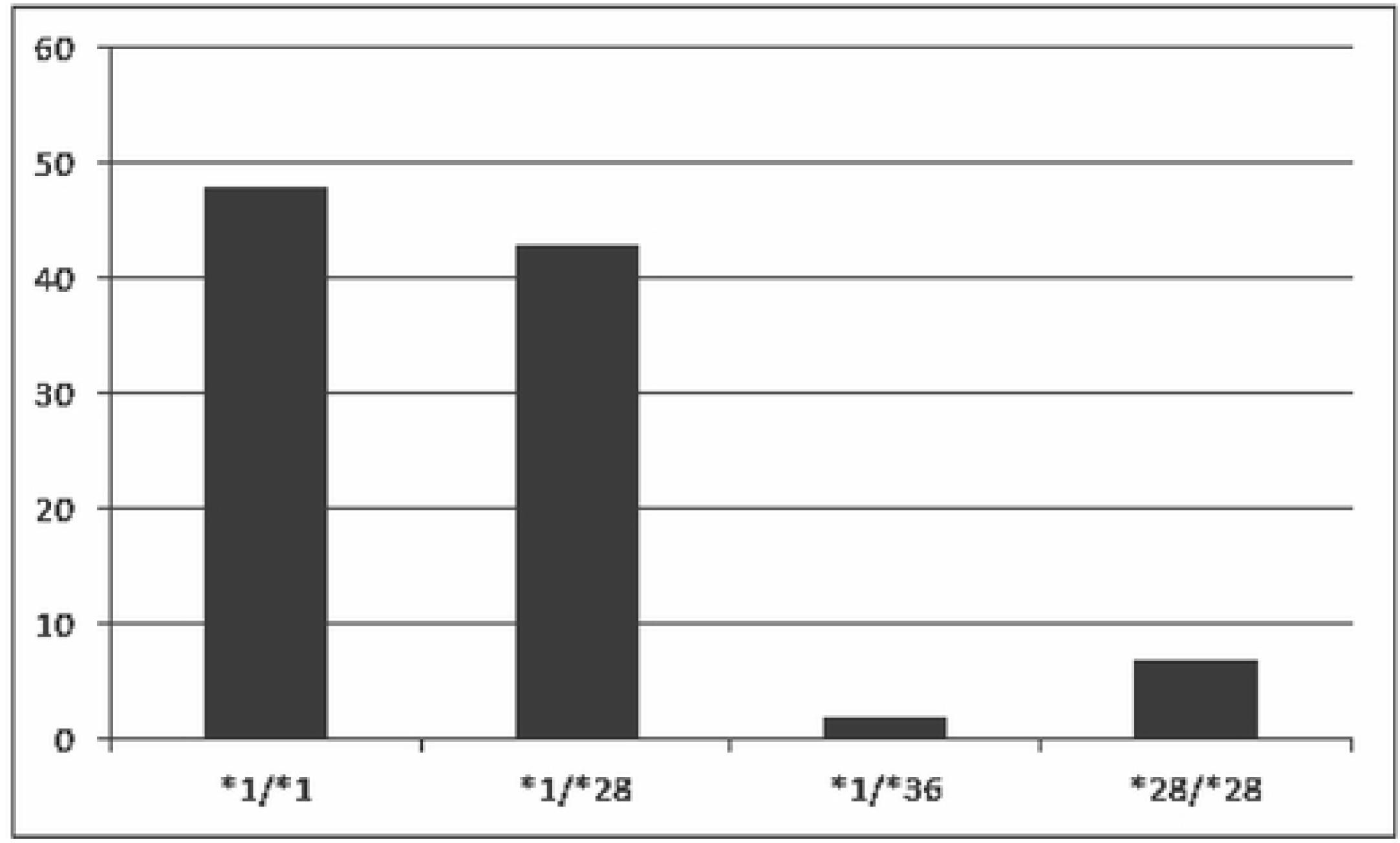
UGT1A1 Genotypes distribution in Argentinian population

## DISCUSSION

UGT 1A1 is the major UGT isoform responsible for glucuronidation of bilirubin in human liver, and it is capable of conjugating various phenols, anthraquinones and flavones. Mutations of bilirubin UDP-glucurunosyl transferase causes hereditary unconjugated hyperbilirubinemia and can also alter the metabolism of certain drugs.

In our population of 100 healthy argentinian volunteers, the distribution of the UGT1A1 showed a predominance of the *1 allele whereas the *28 allele showed a relatively high prevalence of 21,5%.

Compared to this, in a cohort of 245 healthy men and women, aged 20-40 years of Caucasians and Asians the frequencies of the UGT1A1 genotypes were 53,7% for 6/6 *1, 34,8% for 6/7 *1-28, 9,8% for 7/7 *28, 0.8% for 5/6 and 0.8% for 6/8 *1-38 promoter TA repeats. Both allele and genotype frequencies varied by race (P < 0.02), with 11% of the Caucasians and none of the Asians having the 7/7 *28 genotype. Overall, 8% were homozygous variant for both UGT1 polymorphisms and 43% had at least one variant allele for both UGT1A1*28 and UGT1A6*2(11).

The incidences and types of the polymorphisms for these enzymes vary according to geographical regions and ethnic groups. This is the first study that has examined the frequency of polymorphisms for UGT1 isozymes for a population of healthy argentinians.

Since most Argentinian inhabitants are of Caucasian descent, a European genotype frequency profile is to be expected. That is evident in the wild type prevalence in our population and the one from Lampe et. al. However, the contribution of Native American ancestry to gene pool components may in part explain the higher prevalence of the *28 genotype in UGT1A1 *1 in our population, in comparison with the 11% observed in the european study. This finding provides additional information to the ongoing controversy regarding which population -if any- can be used as a general reference for Argentinian genotypic profiles.

A limitation of our results is that all the volunteers evaluated are from the Capital City of Buenos Aires, Argentina. This does leave aside a considerable amount of variant genomics particularly in a vast country that has a longstanding history of racial mixture.

Understanding these polymorphisms and the characterization of these in different populations is essential for the prevention of adverse effects of a considerable number of drugs and also for the reduction of cancer-associated risks.

